# 8-azaadenosine and 8-chloroadenosine are not selective inhibitors of ADAR

**DOI:** 10.1101/2021.05.12.443853

**Authors:** Kyle A. Cottrell, Luisangely Soto Torres, Jason D. Weber

## Abstract

The RNA editing enzyme ADAR, is an attractive therapeutic target for multiple cancers. Through its deaminase activity, ADAR edits adenosine to inosine in dsRNAs. Loss of ADAR in some cancer cell lines causes activation of the type I interferon pathway and the PKR translational repressor, leading to inhibition of proliferation and stimulation of cell death. As such, inhibition of ADAR function is a viable therapeutic strategy for many cancers. However, there are no FDA approved inhibitors of ADAR. Two small molecules have been previously described as inhibitors of ADAR: 8-azaadenosine and 8-chloroadenosine. Here we show that neither molecule is a selective inhibitor of ADAR. Both 8-azaadenosine and 8-chloroadenosine show similar toxicity to ADAR-dependent and independent cancer cell lines. Furthermore, the toxicity of both small molecules is comparable between cell lines with knockdown of ADAR and cells with unperturbed ADAR expression. Treatment with neither molecule causes activation of PKR. Finally, treatment with either molecule has no effect on A-to-I editing of an ADAR substrate. Together these data show that 8-azaadenosine and 8-chloroadenosine are not suitable small molecules for therapies that require selective inhibition of ADAR, and neither should be used in preclinical studies as ADAR inhibitors.

## Introduction

ADAR carries out adenosine-to-inosine (A-to-I) editing within double-stranded RNA (dsRNA) (1-5). By editing dsRNA, it has been proposed that ADAR prevents sensing of self dsRNAs by dsRNA binding proteins involved in activation of the type I interferon (IFN) response and/or control of translation (6-10). Depletion of ADAR in numerous cancer cell lines causes reduced proliferation and increased apoptosis (11-14). Consistent with its proposed role in preventing dsRNA sensing, loss of ADAR in many human cancer cell lines leads to activation of the type I IFN pathway through activation of MAVS and translation repression by activation of PKR (11-13). The growth phenotype of ADAR depletion can be rescued by disruption of type I IFN signaling or knockdown of PKR (11-13). Because of the importance of ADAR expression in many human cancer cell lines, several groups have proposed the use of ADAR inhibitors as a therapy for lung, breast and thyroid cancers (11-14).

There are currently no FDA approved ADAR inhibitors. However, two small molecules have previously been reported to either inhibit ADAR or reduce its expression (14-16). Both of these small molecules are adenosine analogues, Figure 1a. 8-azaadenosine has been used as an ADAR inhibitor in multiple studies involving leukemic stem cells and thyroid cancer cell lines (14,16). In thyroid cancer cell lines, 8-azaadenosine has been shown to be very effective at inhibiting proliferation, even at doses as low as 1-2 µM (14). The use of 8-azaadenosine as an inhibitor of ADAR was initially inspired by a study that incorporated 8-azaadenosine and other adenosine analogues into an ADAR substrate to identify modified substrates that would serve to resolve the structure of ADAR (17). In that study, it was observed that an ADAR substrate containing 8-azadenosine resulted in improved A-to-I editing (17). As such, it is conceivable that free 8-azaadenosine could serve as a competitive inhibitor of ADAR.

**Figure 1:**
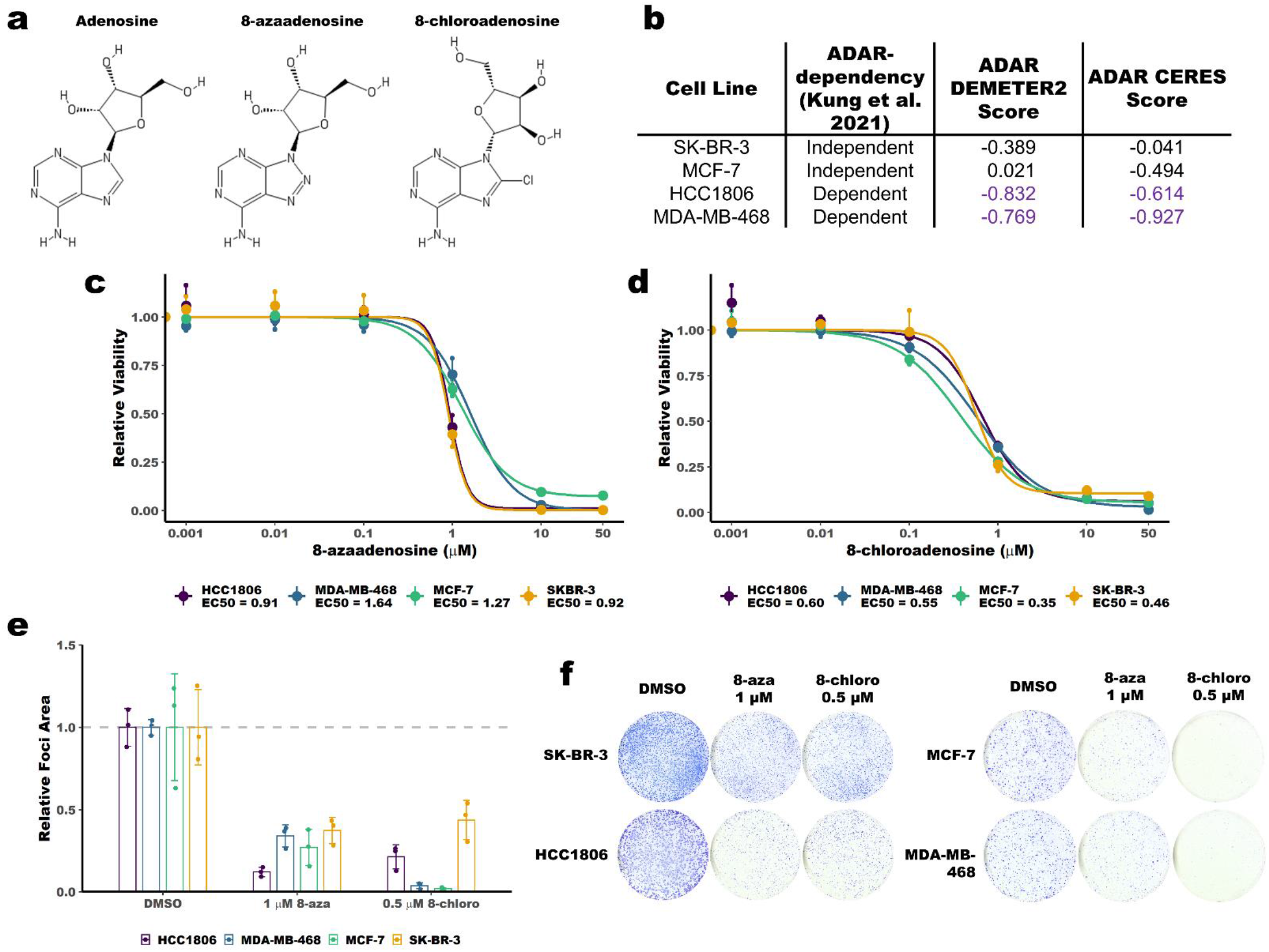
8-chloroadenosine and 8-azaadenosine inhibit proliferation of ADAR-dependent and ADAR-independent breast cancer cell lines. **a** Structure of adenosine, 8-azaadenosine and 8-chloroadenosine. **b** A table summarizing the ADAR-dependency status of relevant breast cancer cell lines as previously published. DEMETER2 corresponds to ADAR-dependency as determined by RNAi screening (20-22). CERES corresponds to ADAR-dependency as determined by CRISPR-Cas9 screening (23,24). A DEMETER2 or CERES score of less than -0.5 is considered “dependent” or “essential” (20,23). **c** Dose response curve for 8-azaadenosine treatment of several breast cancer cell lines. **d** Dose response curve for 8-chloroadenosine treatment of several breast cancer cell lines. In both **c** and **d** cell viability was measured by CellTiter-Glo 2.0. **e** Quantification of foci-formation, panel **f**, following treatment of several breast cancer cell lines with 8-chloroadenosine (8-chloro) or 8-azaadenosine (8-aza). For all panels, error bars are mean +/- standard deviation. In panels **c** and **d**, the large points are the mean of three independent experiments, the smaller points are the mean of three technical replicates performed for each experiment. For panel **e**, the smaller points represent the relative foci area from each of three independent experiments and the column represents the mean foci area of the three experiments.

Another adenosine analogue, 8-chloroadenosine, has been shown not to inhibit the deaminase activity of ADAR itself, but to reduce ADAR expression (15). Treatment of several breast cancer cell lines with 8-chloroadenosine led to reduced ADAR expression and induction of cell death. The cell death phenotype could be rescued by overexpression of wild-type ADAR, but not a dsRNA binding deficient mutant of ADAR, suggesting that 8-chloroadenosine could have some selectivity towards ADAR.

Here we set out to further evaluate the therapeutic potential of 8-chloroadenosine and 8-azaadenosine as ADAR inhibitors. Using several approaches, we show that neither 8-chloroadenosine or 8-azaadenosine are selective inhibitors of ADAR: both molecules inhibit growth of ADAR-depleted cells, treatment with neither molecule caused activation of PKR, and treatment with neither molecule reduced A-to-I editing of an ADAR substrate. Together, these results do not support the use of 8-azaadenosine or 8-chloroadenosine as ADAR inhibitors, and instead warrant the future search for novel ADAR inhibitors.

## Materials and Methods

### Cell culture

Breast cancer cell lines (MCF-7 (RRID:CVCL_0031), SK-BR-3 (RRID:CVCL_0033), HCC1806 (RRID:CVCL_1258), MDA-MB-468 (RRID:CVCL_0419)) were obtained from American Type Culture Collection, in 2011. All cell lines were cultured in Dulbecco’s modified Eagle’s medium (DMEM) (Hyclone) with 10% fetal bovine serum (Invitrogen), 2 mM glutamine (Hyclone), 0.1 mM nonessential amino acids (Hyclone), 1 mM sodium pyruvate (Hyclone), and 2 μg/ml gentamicin (Invitrogen). 8-chloroadenosine and 8-azaadenosine were purchased from Tocis, catalogue numbers: 4436 and 6868.

### Viral Production and Transduction

Lentivirus was produced by Turbo DNAfectin 3000 (Lambda Biotech) transfection of 293T cells with pCMV-VSV-G, pCMV-ΔR8.2, and pLKO.1-puro for shRNAs. Virus was harvested 48 hours post-transfection. Cells were transduced with lentivirus for 16 hours in the presence of 10 µg/mL protamine sulfate. The cells were selected with puromycin at 2 µg/mL for one day. For analysis of ADAR expression and PKR activation following ADAR knockdown, cells were harvested 96 hours after transduction. The sequences for the shRNA-scramble (shSCR) and shADAR were described and validated previously (13).

### Data and Code Availability

Scripts used for all plots are available on GitHub (https://github.com/cottrellka/ADAR_5-2021).

### Immunoblot

Cell pellets were lysed and sonicated in RIPA Buffer (50 mM Tris pH 7.4, 150 mM NaCl, 1% Triton X-100, 0.1% sodium dodecyl sulfate and 0.5% sodium deoxycholate) with 1x HALT Protease Inhibitor (Pierce). Forty micrograms of protein lysate were resolved on 4-12% TGX Acrylamide Stain-Free gels (Bio-Rad). Proteins were transferred to PVDF membrane (Millipore). The membrane was cut into strips corresponding to the molecular weight of proteins of interest. The blots were blocked and then probed with the appropriate primary antibodies: Primary antibodies: ADAR1 (Santa Cruz, sc-73408), PKR (Cell Signaling, #3072), PKR Thr-446-P (Abcam, ab32036), GAPDH (Bethyl, A300-641A). Primary antibodies were detected with horseradish-peroxidase conjugated secondary antibodies (Jackson ImmunoResearch) and detection was carried out with Clarity Western ECL Substrate (Bio-Rad). Chemiluminescence was imaged using a ChemiDoc imaging system (Bio-Rad). Quantification of immunoblots was performed using Image Lab software (Bio-Rad). All proteins were normalized to GAPDH abundance. For PKR and pPKR, two separate gels were resolved, transferred, and probed for either PKR or pPKR in addition to GAPDH for both. PKR and pPKR abundance were normalized to GAPDH prior to normalizing pPKR to PKR. Uncropped immunoblot images are available in Supplemental Figures 1-5.

### Analysis of A-to-I Editing

Cells were treated as indicated for 72 hours prior to harvesting of RNA using the Nucleospin RNA kit (Macherey-Nagel). First-strand cDNA synthesis was performed using iScript Supermix (Bio-Rad). The cDNA was purified using the Monarch DNA and PCR Cleanup Kit (New England Biolabs). A region around an A-to-I editing site in BPNT1 was amplified by Q5 Hot Start High-Fidelity 2X Master Mix (New England Biolabs) and the primers: BPNT1_F 5’-TGCTGTGGGAGGCAAGTTAAC-3’ and BPNT1_R 5’-GAGTCCGAGGCAGACAGATC-3’. The PCR parameters were as follows: 98 °C for 30 s, 98 °C for 30 s, 72 °C for 30 s, 72 °C for 55 s, repeat steps 2-4 for 19 cycles dropping the annealing temperature 0.2 °C each cycle, 98 °C for 30 s, 68 °C for 30 s, 72 °C for 55 s, repeat steps 6-8 for 19 cycles, 72 °C for 5 minutes. The PCR products were resolved by agarose gel electrophoresis and purified using the Monarch Gel Extraction kit (New England Biolabs). Purified PCR products were Sanger sequenced by Genewiz using the BPNT1_F_Seq primer: 5’-GGAGTCTCGCTCTGTAGCCT-3’. The chromatograms for all replicates are available in Supplemental Figures 5-8. To determine percent editing, raw peak heights were measured for the edited and unedited base using the program QSVanalyzer (18). Percent editing was calculated by the following formula:

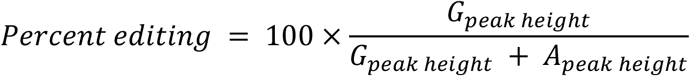

### Measurement of Cell Viability

Cells were treated as indicated for 96 hours prior to assessment of cell viability using CellTiter-Glo 2.0 (Promega) per manufacturers’ protocol. Luminescence was measured for 10 s using a Promega Glomax Navigator luminometer. Dose response analysis was performed using the R package ‘drc’ (19). A four-parameter log-logistic model (LL.4) was fit to the viability data. For this log-logistic model, the Hill Coefficient, lower limit, and EC50 were allowed to vary but the upper limit was set to 1. Further details for this analysis can be found in the GitHub repository above.

### Foci Formation Assay

Five thousand cells were plated for each condition in a 10 cm culture dish. Three days later the cells were treated as indicated. After 9 (HCC1806 and SK-BR-3) to 14 (MCF-7 and MDA-MB-468) days the cells were washed briefly with 1x PBS prior to fixation in 100% methanol. After drying, the cells were stained with Giemmsa (Sigma-Aldrich) prior to washing excess stain away with deionized water. The plates were scanned using an ImageScanner III (General Electric). Foci area was calculated using ImageJ.

## Results

### Cytotoxicty 8-chloroadenosine and 8-azaadenosine in breast cancer cell lines

Knockdown or knockout of ADAR causes reduced proliferation and increased cell death in numerous, but not all cancer cell lines (11-14). ADAR-dependency has been evaluated through large screening experiments (20-24) and smaller studies involving knockdown or knockout of ADAR in panels of human cancer cell lines (11-14). Recently, ADAR-dependency was evaluated for a panel of human breast cancer cell lines (13). To evaluate the on-target effects of 8-chloroadenosine and 8-azaadenosine, we assessed the effects of each small molecule on cell viability of breast cancer cell lines previously identified to be ADAR-dependent or -independent, Figure 1b. If 8-chloroadenosine and/or 8-azaadenosine are selective inhibitors of ADAR, it would be expected that the EC50 for cell viability of each drug would be lower for ADAR-dependent cell lines relative to ADAR-independent cell lines. However, analysis of the effects of each adenosine analogue on cell viability found that the EC50s were comparable between ADAR-dependent and independent cell lines, Figure 1c-d. For 8-chloroadenosine there was a ∼0.25 µM EC50 difference between the most sensitive cell line (MCF-7, ADAR-independent) and the least (HCC1806, ADAR-dependent). Similarly, for 8-azaadenosine there was a < 1 µM EC50 difference between the most sensitive cell line (SK-BR-3, ADAR-independent) and least sensitive (MDA-MB-468, ADAR-dependent). These data were largely supported by foci formation analysis, Figure 1e-f. The ADAR-independent cell lines SK-BR-3 and MCF-7, and the ADAR-dependent cell line MDA-MB-468 were similarly sensitive to the effects of 8-azadenosine on foci formation. The two cell lines most sensitive to the effects of 8-chloroadenosine on foci formation were MCF-7 and MDA-MB-468, ADAR-independent and ADAR-dependent cell lines, respectively. Taken together, these data show that neither 8-chloroadenosine or 8-azaadenosone are selectively cytotoxic towards ADAR-dependent cell lines.

### Cytotoxicity of 8-chloroadenosine and 8-azadenosine in ADAR-depleted cells

While the data described in Figure 1 are consistent with 8-azaadenosine and 8-chloradenosine lacking selectivity for ADAR, we sought to address this question more thoroughly by assessing the cytotoxicity of the small-molecules in ADAR-depleted cell lines. ADAR was knocked-down in two ADAR-independent cell lines, SK-BR-3 and MCF-7, Figure 2a and 2d. The EC50 of cell viability for 8-azaadenosine and 8-chloroadeonsine was evaluated for control (shSCR) or ADAR knockdown (shADAR). If 8-azaadenosine and/or 8-chloroadenosine are selective inhibitors of ADAR, it would be expected that ADAR-depleted cells would be less sensitive to each adenosine analogues. However, the EC50 for each drug was generally similar between shSCR and shADAR transduced cells for both cell lines, Figure 2b-c and 2e-f. Only for 8-chloroadenosine was there a clear difference between the EC50 in shSCR versus shADAR transduced cells, with shADAR cells having a lower EC50. Together these data, and the data in Figure 1, show that neither 8-chloroadenosine or 8-azaadenosone induce cytotoxicity through selective inhibition of ADAR.

**Figure 2:**
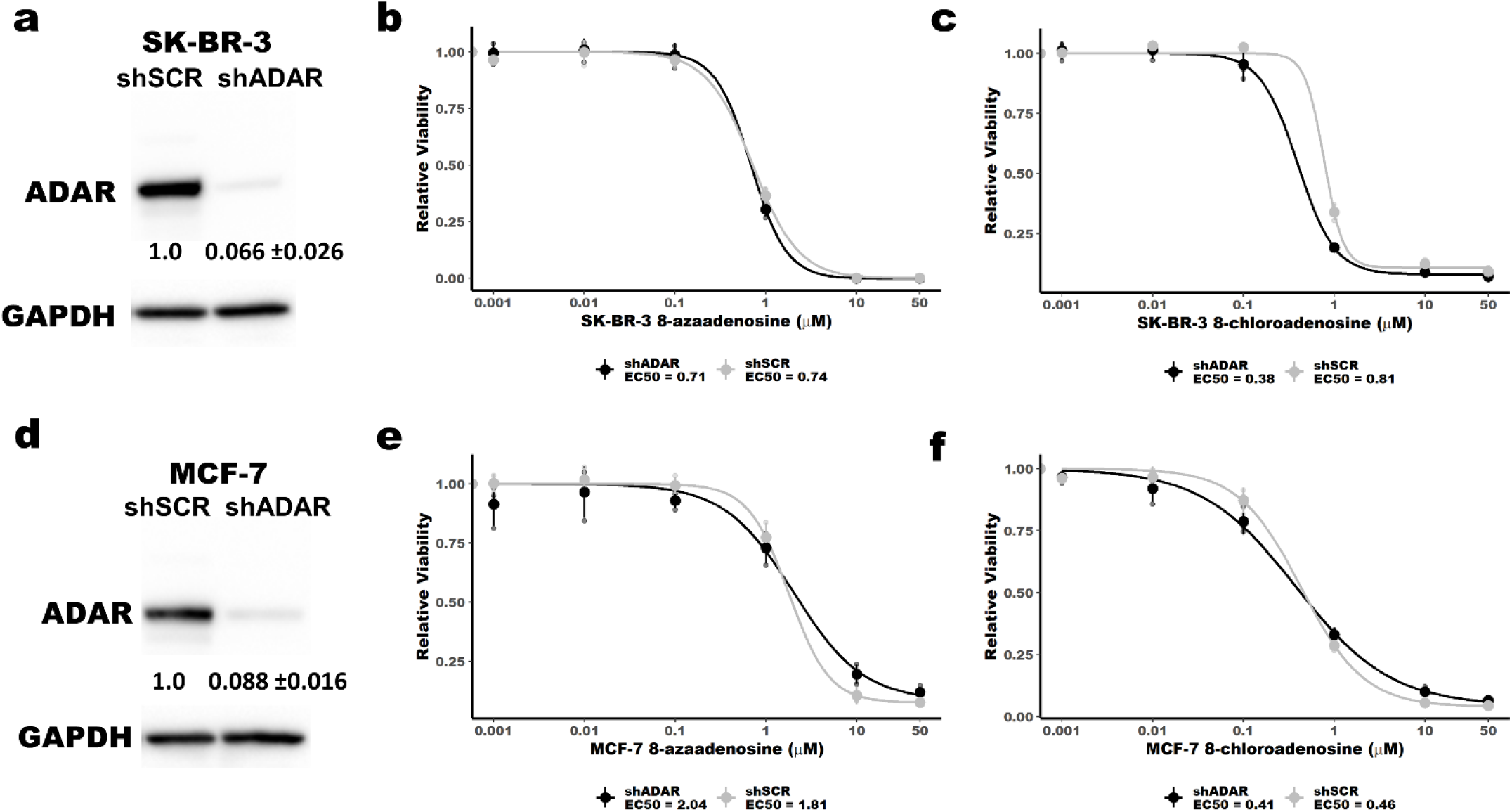
8-chloroadenosine and 8-azaadenosine inhibit proliferation of ADAR depleted breast cancer cell lines. Immunoblot of ADAR knockdown in SK-BR-3 (**a**) and MCF-7 (**d**). The level of ADAR knockdown is shown below each band, mean +/- standard deviation. Five (SK-BR-3) or six (MCF-7) days after transduction of shSCR or shADAR, the cells were treated with 8-chloroadenosine or 8-azaadenosine for dose response curves. **b** and **c**, Dose response curves for 8-azaadenosine and 8-chloroadenosine in SK-BR-3 cells with (shADAR) or without (shSCR) ADAR knockdown. **e** and **f**, Dose response curves for 8-azaadenosine and 8-chloroadenosine in MCF-7 cells with (shADAR) or without (shSCR) ADAR knockdown. In panels **b, c, e**, and **f** the large points are the mean of three independent experiments, the smaller points are the mean of three technical replicates performed for each experiment, error bars are mean +/- standard deviation.

### Treatment with 8-chloroadenosine or 8-azaadenosine does not activate PKR

Loss of ADAR in ADAR-dependent cells has been shown to cause activation of the dsRNA sensor PKR (11-13). It has been proposed that loss of A-to-I editing by ADAR leads to accumulation of dsRNA leading to activation and autophosphorylation of PKR (9). Activation of PKR leads to inhibition of translation and induction of cell death (11-13). Selective inhibitors of ADAR would be expected to also cause PKR activation. We evaluated PKR activation upon treatment with 8-chloroadenosine or 8-azadenosine by immunoblot using a phospho-PKR (phospho-T446) specific antibody. Unlike knockdown of ADAR, which caused robust activation of PKR in the ADAR-dependent cell line HCC1806 and MDA-MB-468, Figure 3a and 3c-d, neither 8-chloroadenosine or 8-azaadenosine induced PKR activation in the same cell lines, Figure 3b, 3e-i. These data suggest that neither 8-chloroadenosine or 8-azadenosine are inhibitors of ADAR.

**Figure 3:**
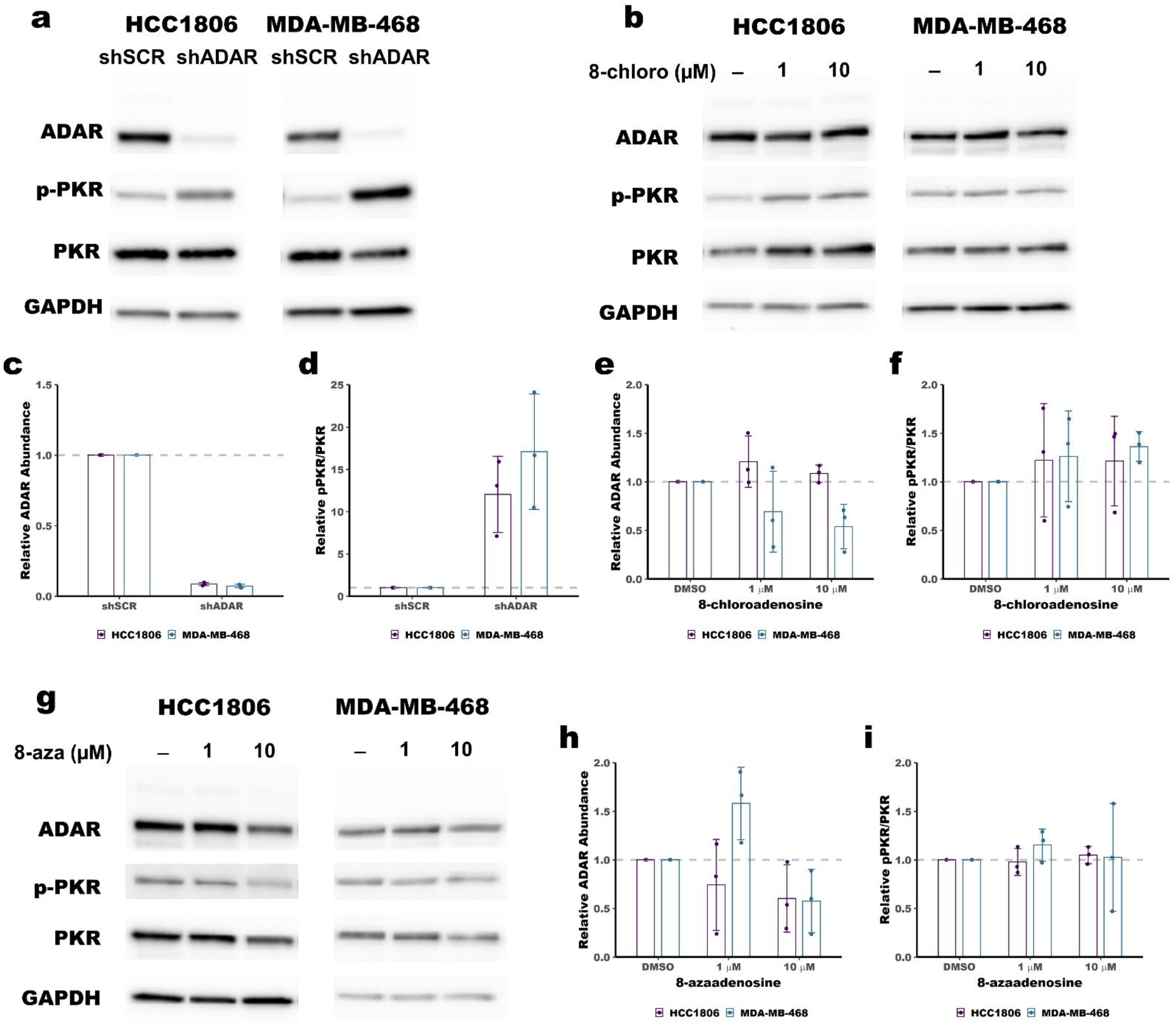
Treatment with 8-chloroadenosine or 8-azaadenosine does not activate PKR. **a** Immunoblot showing activation of PKR (increased phosphorylation of PKR at T446, pPKR) following knockdown of ADAR in HCC1806 and MDA-MB-468. **c** and **d** quantification of the immunoblot in panel **a. b** Immunoblot showing no activation of PKR following treatment of HCC1806 and MDA-MB-468 with 8-chloroadenosine (8-chloro). **e** and **f** quantification of the immunoblot in panel **b. g** Immunoblot showing no activation of PKR following treatment of HCC1806 and MDA-MB-468 with 8-azaadenosine (8-aza). **h** and **i** quantification of the immunoblot in panel **g**. For panel **c-f** and **h-i**, the smaller points represent relative ADAR abundance or relative pPKR/PKR from each of three independent experiments, and the column represents the mean of the three experiments. Error bars are mean +/- standard deviation.

### Treatment with 8-chloroadenosine or 8-azaadenosine has no effect on A-to-I editing

To directly test the effects of 8-azaadenoinse and 8-chloroadenosine on the deaminase activity of ADAR, we used Sanger sequencing to measure A-to-I editing of a highly edited ADAR substrate – BPNT1 (25). The adenosine at position 1894 in the BPNT1 mRNA was shown to be highly edited ∼75% in four different breast cancer cell lines (25). Percent editing can be measured by Sanger sequencing of PCR amplified cDNA. As inosine pairs most readily with cytosine, reverse transcriptase will incorporate a cytosine at each A-to-I editing event. Sanger sequencing of the PCR product made from the cDNA will show either an A (for unedited transcripts) or a G (for edited transcripts). We performed this analysis to assess the change in A-to-I editing of BPNT1-A1894 upon ADAR knockdown. Knockdown of ADAR reduced editing by ∼3-fold, Figure 4a-b. The same analysis was performed for cells treated with either 1 or 10 µM 8-azaadenosone or 8-chloroadenosine. There were no substantial changes to editing of BPNT1-A1894 upon treatment with either adenosine analogue, Figure 4c-f. Together, these data clearly show that neither 8-chloroadenosine or 8-azaadenosine affects A-to-I editing of BPNT1-A1894.

**Figure 4:**
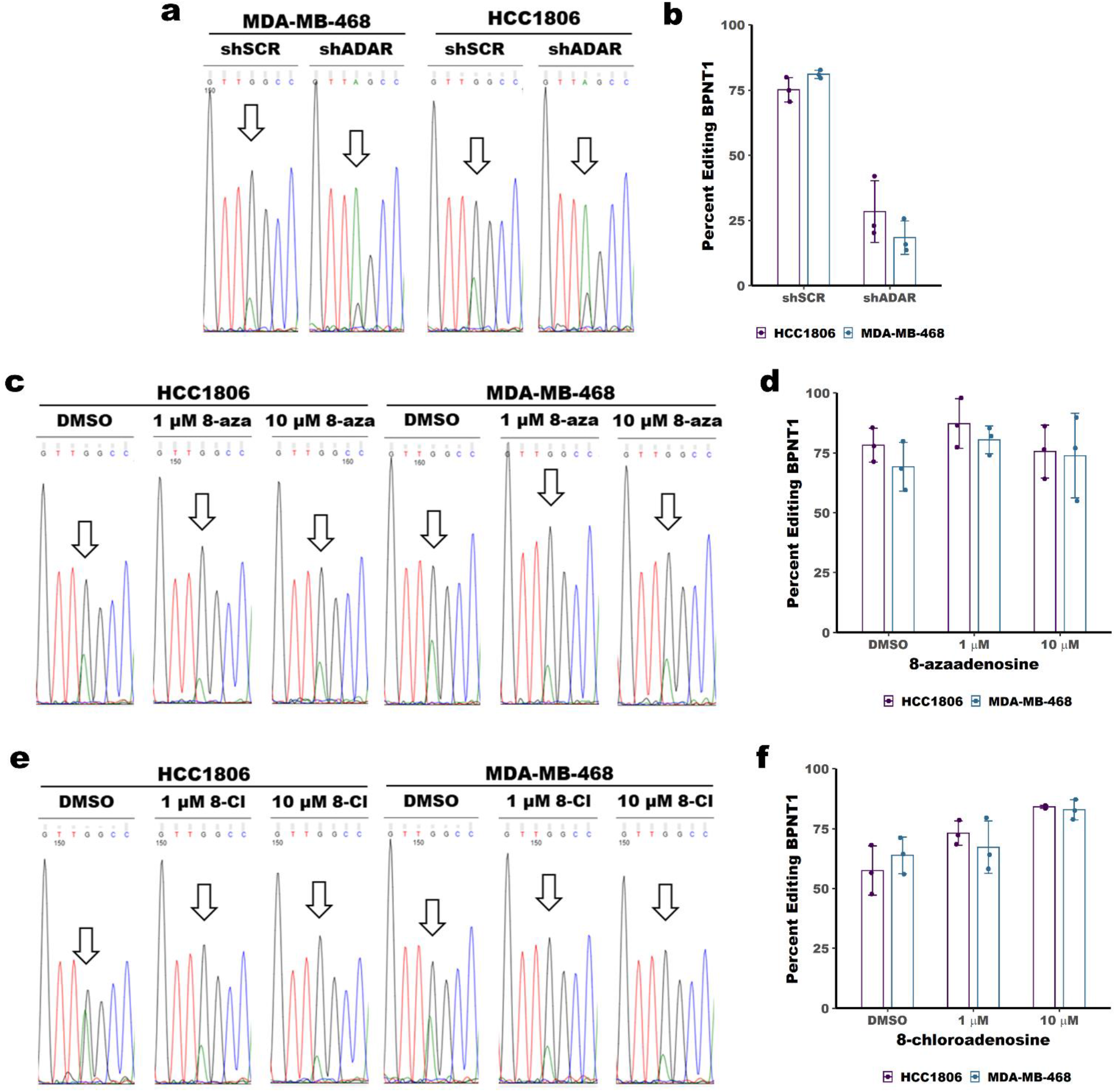
Treatment with 8-chloroadenosine or 8-azaadenosine does not affect A-to-I editing. **a** Sanger sequencing chromatogram of BPNT1 with or without ADAR knockdown. The arrow indicates a base edited by ADAR. The editing site is at position 1894 within the BPNT1 transcript (NM_006085.6). **b** Quantification of percent editing as measured by Sanger sequencing in panel **a**. Percent editing was calculated as the edited base (G) peak height divided by the total peak height of the unedited (A) and edited (G) base. **c** Sanger sequencing chromatogram of BPNT1 with or without 8-azaadenosine (8-aza) treatment. **d** Quantification of editing efficiency from panel **c. e** Sanger sequencing chromatogram of BPNT1 with or without 8-azaadenosine (8-aza) treatment. **f** Quantification of editing efficiency from panel **e**. For panels **b, d** and **f**, the smaller points represent percent editing from each of three independent experiments, and the column represents the mean of the three experiments. Error bars are mean +/- standard deviation.

## Discussion

Several recent studies have highlighted the importance of ADAR expression in a wide range of cancer cell lines (11-14). In ADAR-dependent cells, loss of ADAR causes activation of PKR and the type I IFN pathway leading to reduced proliferation and apoptosis. Furthermore, loss of ADAR in cell lines that do not require ADAR expression to grow in tissue culture conditions has been shown to improve anti-tumor immunity *in vivo*, especially in combination with anti-PD1 therapies (26). The importance of ADAR in tumor biology therefore makes it an ideal therapeutic target for multiple cancers.

While there are currently no FDA approved ADAR inhibitors available for clinical use, two adenosine analogues have been used in pre-clinical studies to perturb ADAR activity or expression – 8-chloroadenosine and 8-azaadenosine. We found that both adenosine analogues efficiently reduce the viability of both ADAR-dependent and ADAR-independent cell lines. Similarly, both adenosine analogues reduced the viability of ADAR-depleted cell lines to a similar or greater extent than cell lines with unperturbed ADAR expression. We showed that treatment with neither 8-chloradenosine or 8-azaadenosine caused activation of PKR, in contrast with ADAR-knockdown which caused robust PKR activation in the same cell lines. Finally, we observed that neither adenosine analogue inhibited A-to-I editing of an ADAR substrate.

The off-target effects of either 8-chloroadenosine or 8-azaadenosine are consistent with what is known about the biological activity of both adenosine analogues. It has been shown that both adenosine analogues can be incorporated into nascent RNA and DNA (27-29), and both have been shown to inhibit DNA synthesis (28,30). Furthermore, both 8-azaadenosine and 8-chloroadenosine can be rapidly incorporated into the cellular ATP pool, replacing ATP with 8-azaATP or 8-chloroATP (29-32). 8-chloadenosine has also been shown to cause inhibition of mTOR and activation of AMPK in renal cell carcinoma cell lines (33). Additionally, 8-chloradenosine has been shown to activate the unfolded protein response leading to apoptosis in coronary artery endothelial cells (29). Finally, *in vivo* studies of 8-azaadenosine toxicity revealed significant hepatic toxicity (31). Taken together, these previous findings, along with those presented here, show that 8-chloradenosine or 8-azaadenosine likely cause cell death through numerous indirect effects and not through selective inhibition of ADAR. Neither 8-azaadenosine or 8-chloroadenosine should be used as ADAR inhibitors.

## Acknowledgements

This work was supported by NIH grant R01CA190986 and Department of Defense Breast Cancer Research Program grant W81XWH-18-1-0025. This work was supported by the Longer Life Foundation: A RGA/Washington University partnership.

## Author Contributions

KAC conceived the project. KAC and LST performed the experiments and data analysis. KAC wrote the manuscript. All authors edited the manuscript.

## Competing interests

The author(s) declare no competing interests.

## Notes

Conflict of interest statement: JDW is a compensated consultant for Ono Pharmaceuticals. The other authors declare no potential conflicts of interest.

### Competing Interest Statement

JDW is a compensated consultant for Ono Pharmaceuticals.
The other authors have no competing interest.

## References

1. Cattenoz PB, Taft RJ, Westhof E, Mattick JS. Transcriptome-wide identification of A > I RNA editing sites by inosine specific cleavage. RNA 2013;19(2):257–70 doi 10.1261/rna.036202.112.

2. Carmi S, Borukhov I, Levanon EY. Identification of widespread ultra-edited human RNAs. PLoS Genet 2011;7(10):e1002317 doi 10.1371/journal.pgen.1002317.

3. Morse DP. Identification of substrates for adenosine deaminases that act on RNA. Methods Mol Biol 2004;265:199–218 doi 10.1385/1-59259-775-0:199.

4. Bazak L, Levanon EY, Eisenberg E. Genome-wide analysis of Alu editability. Nucleic Acids Res 2014;42(11):6876–84 doi 10.1093/nar/gku414.

5. Bazak L, Haviv A, Barak M, Jacob-Hirsch J, Deng P, Zhang R, et al. A-to-I RNA editing occurs at over a hundred million genomic sites, located in a majority of human genes. Genome Res 2014;24(3):365–76 doi 10.1101/gr.164749.113.

6. Liddicoat BJ, Piskol R, Chalk AM, Ramaswami G, Higuchi M, Hartner JC, et al. RNA editing by ADAR1 prevents MDA5 sensing of endogenous dsRNA as nonself. Science 2015;349(6252):1115–20 doi 10.1126/science.aac7049.

7. George CX, Ramaswami G, Li JB, Samuel CE. Editing of Cellular Self-RNAs by Adenosine Deaminase ADAR1 Suppresses Innate Immune Stress Responses. The Journal of biological chemistry 2016;291(12):6158–68 doi 10.1074/jbc.M115.709014.

8. Lamers MM, van den Hoogen BG, Haagmans BL. ADAR1: “Editor-in-Chief” of Cytoplasmic Innate Immunity. Front Immunol 2019;10:1763 doi 10.3389/fimmu.2019.01763.

9. Chung H, Calis JJA, Wu X, Sun T, Yu Y, Sarbanes SL, et al. Human ADAR1 Prevents Endogenous RNA from Triggering Translational Shutdown. Cell 2018;172(4):811-24.e14 doi 10.1016/j.cell.2017.12.038.

10. Pestal K, Funk CC, Snyder JM, Price ND, Treuting PM, Stetson DB. Isoforms of RNA-Editing Enzyme ADAR1 Independently Control Nucleic Acid Sensor MDA5-Driven Autoimmunity and Multi-organ Development. Immunity 2015;43(5):933–44 doi 10.1016/j.immuni.2015.11.001.

11. Liu H, Golji J, Brodeur LK, Chung FS, Chen JT, deBeaumont RS, et al. Tumor-derived IFN triggers chronic pathway agonism and sensitivity to ADAR loss. Nat Med 2019;25(1):95–102 doi 10.1038/s41591-018-0302-5.

12. Gannon HS, Zou T, Kiessling MK, Gao GF, Cai D, Choi PS, et al. Identification of ADAR1 adenosine deaminase dependency in a subset of cancer cells. Nat Commun 2018;9(1):5450 doi 10.1038/s41467-018-07824-4.

13. Kung CP, Cottrell KA, Ryu S, Bramel ER, Kladney RD, Bao EA, et al. Evaluating the therapeutic potential of ADAR1 inhibition for triple-negative breast cancer. Oncogene 2021;40(1):189–202 doi 10.1038/s41388-020-01515-5.

14. Ramírez-Moya J, Baker AR, Slack FJ, Santisteban P. ADAR1-mediated RNA editing is a novel oncogenic process in thyroid cancer and regulates miR-200 activity. Oncogene 2020;39(18):3738–53 doi 10.1038/s41388-020-1248-x.

15. Ding HY, Yang WY, Zhang LH, Li L, Xie F, Li HY, et al. 8-Chloro-Adenosine Inhibits Proliferation of MDA-MB-231 and SK-BR-3 Breast Cancer Cells by Regulating ADAR1/p53 Signaling Pathway. Cell Transplant 2020;29:963689720958656 doi 10.1177/0963689720958656.

16. Zipeto MA, Court AC, Sadarangani A, Delos Santos NP, Balaian L, Chun HJ, et al. ADAR1 Activation Drives Leukemia Stem Cell Self-Renewal by Impairing Let-7 Biogenesis. Cell Stem Cell 2016;19(2):177–91 doi 10.1016/j.stem.2016.05.004.

17. Véliz EA, Easterwood LM, Beal PA. Substrate analogues for an RNA-editing adenosine deaminase: mechanistic investigation and inhibitor design. J Am Chem Soc 2003;125(36):10867–76 doi 10.1021/ja029742d.

18. Carr IM, Robinson JI, Dimitriou R, Markham AF, Morgan AW, Bonthron DT. Inferring relative proportions of DNA variants from sequencing electropherograms. Bioinformatics 2009;25(24):3244–50 doi 10.1093/bioinformatics/btp583.

19. Ritz C, Baty F, Streibig JC, Gerhard D. Dose-Response Analysis Using R. PLoS One 2015;10(12):e0146021 doi 10.1371/journal.pone.0146021.

20. McFarland JM, Ho ZV, Kugener G, Dempster JM, Montgomery PG, Bryan JG, et al. Improved estimation of cancer dependencies from large-scale RNAi screens using model-based normalization and data integration. Nat Commun 2018;9(1):4610 doi 10.1038/s41467-018-06916-5.

21. Marcotte R, Sayad A, Brown KR, Sanchez-Garcia F, Reimand J, Haider M, et al. Functional Genomic Landscape of Human Breast Cancer Drivers, Vulnerabilities, and Resistance. Cell 2016;164(1-2):293–309 doi 10.1016/j.cell.2015.11.062.

22. McDonald ER, de Weck A, Schlabach MR, Billy E, Mavrakis KJ, Hoffman GR, et al. Project DRIVE: A Compendium of Cancer Dependencies and Synthetic Lethal Relationships Uncovered by Large-Scale, Deep RNAi Screening. Cell 2017;170(3):577-92.e10 doi 10.1016/j.cell.2017.07.005.

23. Meyers RM, Bryan JG, McFarland JM, Weir BA, Sizemore AE, Xu H, et al. Computational correction of copy number effect improves specificity of CRISPR-Cas9 essentiality screens in cancer cells. Nat Genet 2017;49(12):1779–84 doi 10.1038/ng.3984.

24. Dempster JM, Rossen J, Kazachkova M, Pan J, Kugener G, Root DE, et al. Extracting Biological Insights from the Project Achilles Genome-Scale CRISPR Screens in Cancer Cell Lines. bioRxiv 2019:720243 doi 10.1101/720243.

25. Fumagalli D, Gacquer D, Rothé F, Lefort A, Libert F, Brown D, et al. Principles Governing A-to-I RNA Editing in the Breast Cancer Transcriptome. Cell Rep 2015;13(2):277–89 doi 10.1016/j.celrep.2015.09.032.

26. Ishizuka JJ, Manguso RT, Cheruiyot CK, Bi K, Panda A, Iracheta-Vellve A, et al. Loss of ADAR1 in tumours overcomes resistance to immune checkpoint blockade. Nature 2019;565(7737):43–8 doi 10.1038/s41586-018-0768-9.

27. Bennett LL, Allan PW. Metabolism and metabolic effects of 8-azainosine and 8-azaadenosine. Cancer Res 1976;36(11 Pt 1):3917–23.

28. Glazer RI, Lloyd LS. Effects of 8-azaadenosine and formycin on cell lethality and the synthesis and methylation of nucleic acids in human colon carcinoma cells in culture. Biochem Pharmacol 1982;31(20):3207–14 doi 10.1016/0006-2952(82)90551-2.

29. Tang V, Fu S, Rayner BS, Hawkins CL. 8-Chloroadenosine induces apoptosis in human coronary artery endothelial cells through the activation of the unfolded protein response. Redox Biol 2019;26:101274 doi 10.1016/j.redox.2019.101274.

30. Dennison JB, Balakrishnan K, Gandhi V. Preclinical activity of 8-chloroadenosine with mantle cell lymphoma: roles of energy depletion and inhibition of DNA and RNA synthesis. Br J Haematol 2009;147(3):297–307 doi 10.1111/j.1365-2141.2009.07850.x.

31. Spremulli EN, Crabtree GW, Dexter DL, Chu SH, Farineau DM, Ghoda LY, et al. Biochemical pharmacology and toxicology of 8-azaadenosine alone and in combination with 2’-deoxycoformycin (pentostatin). Biochem Pharmacol 1982;31(14):2415–21 doi 10.1016/0006-2952(82)90538-x.

32. Macer-Wright JL, Sileikaite I, Rayner BS, Hawkins CL. 8-Chloroadenosine Alters the Metabolic Profile and Downregulates Antioxidant and DNA Damage Repair Pathways in Macrophages. Chem Res Toxicol 2020;33(2):402–13 doi 10.1021/acs.chemrestox.9b00334.

33. Kearney AY, Fan YH, Giri U, Saigal B, Gandhi V, Heymach JV, et al. 8-Chloroadenosine Sensitivity in Renal Cell Carcinoma Is Associated with AMPK Activation and mTOR Pathway Inhibition. PLoS One 2015;10(8):e0135962 doi 10.1371/journal.pone.0135962.

