## Supplemental Information for "8-azaadenosine and 8-chloroadenosine are not selective inhibitors of ADAR"

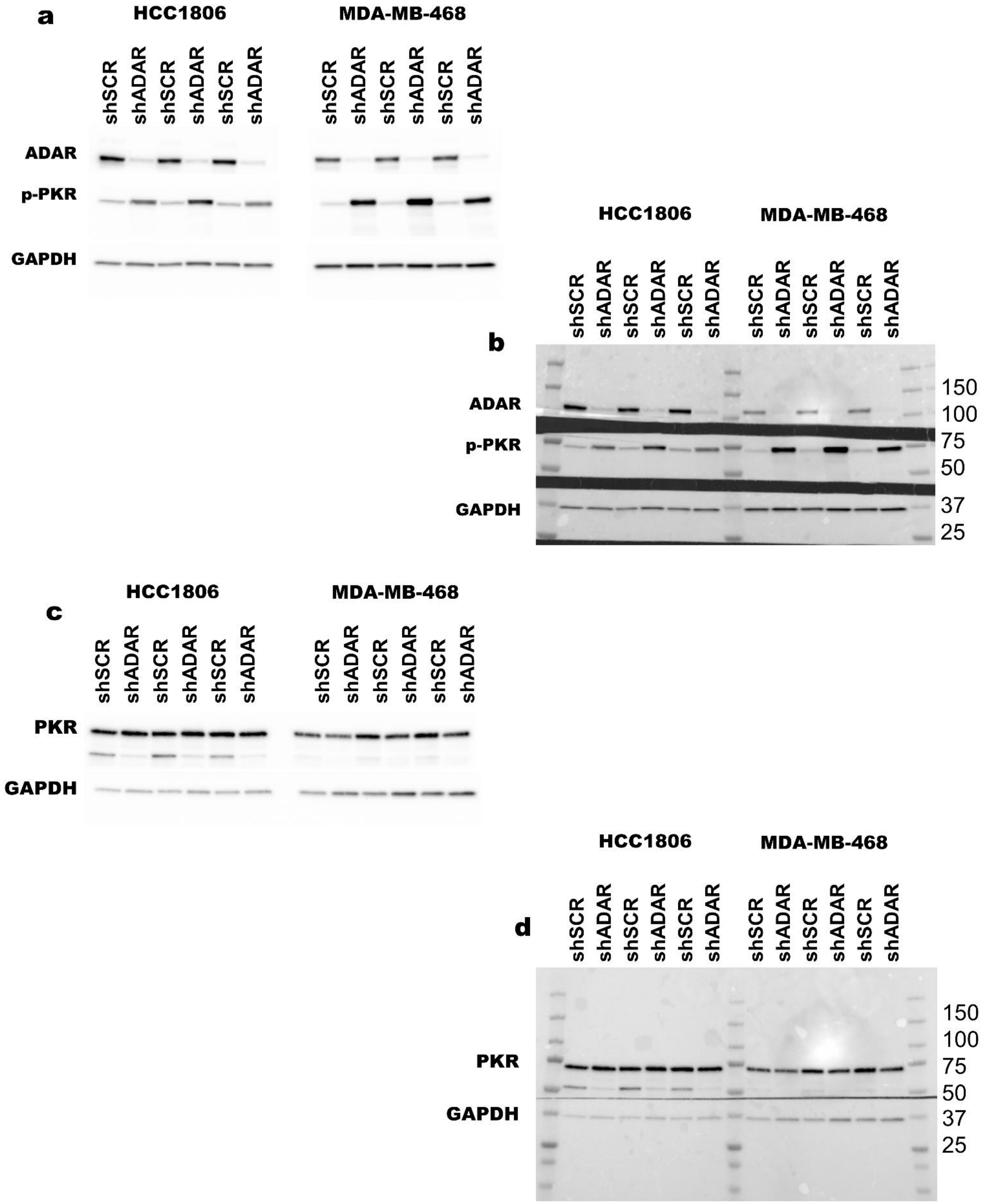


**Supplemental Figure 1:**

Uncropped immunoblots associated with Figure 3A, 3C and 3D. Panels **a** and **b** are the uncropped chemiluminescence images, panels **b** and **d** are the chemiluminescence images merged with colorimetric images to show the molecular weight marker.

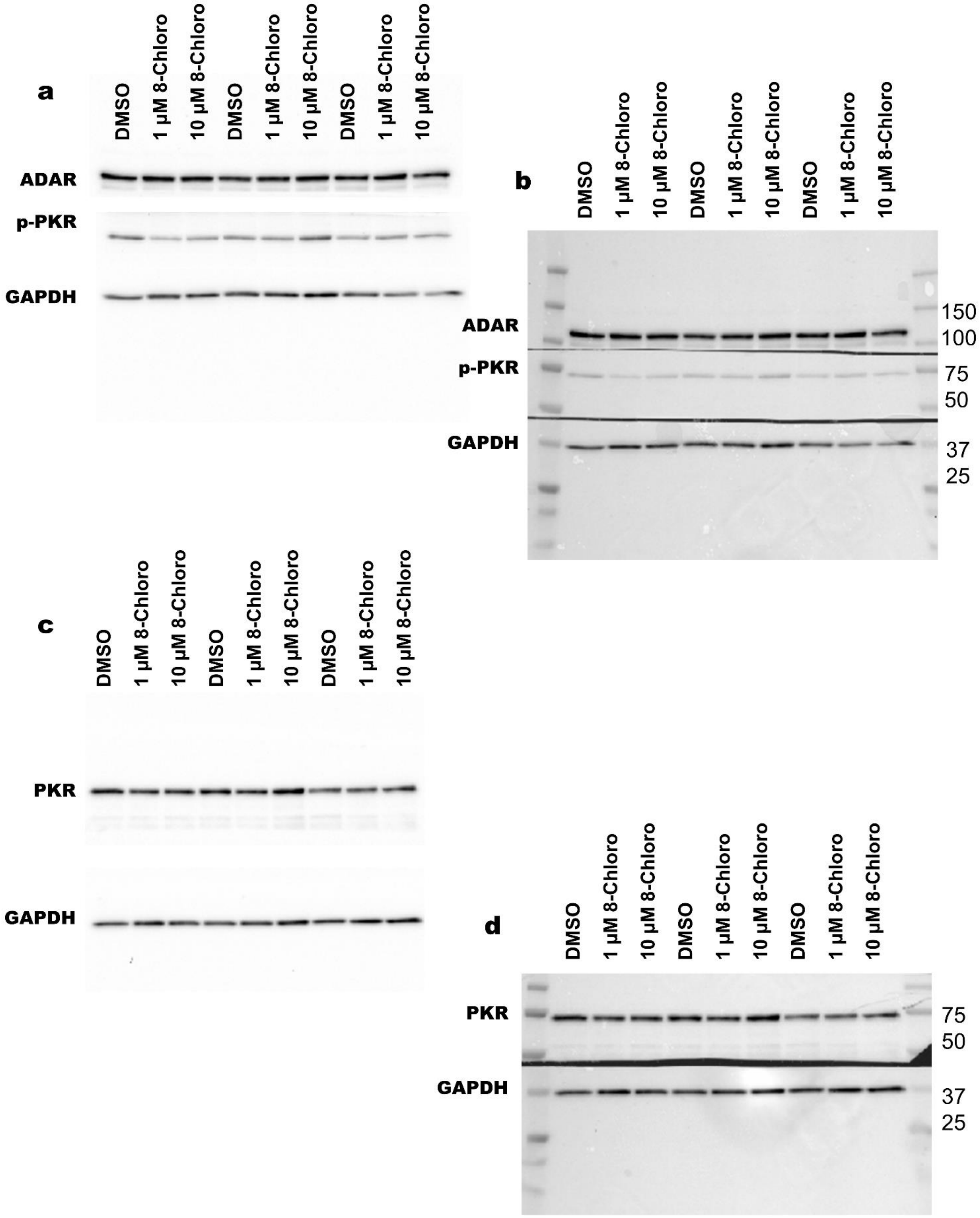


**Supplemental Figure 2:**

Uncropped immunoblots for MDA-MB-468 treatment with 8-chloroadenosine (8-chloro) associated with Figure 3b, 3e and 3f. Panels **a** and **b** are the uncropped chemiluminescence images, panels **b** and **d** are the chemiluminescence images merged with colorimetric images to show the molecular weight marker.

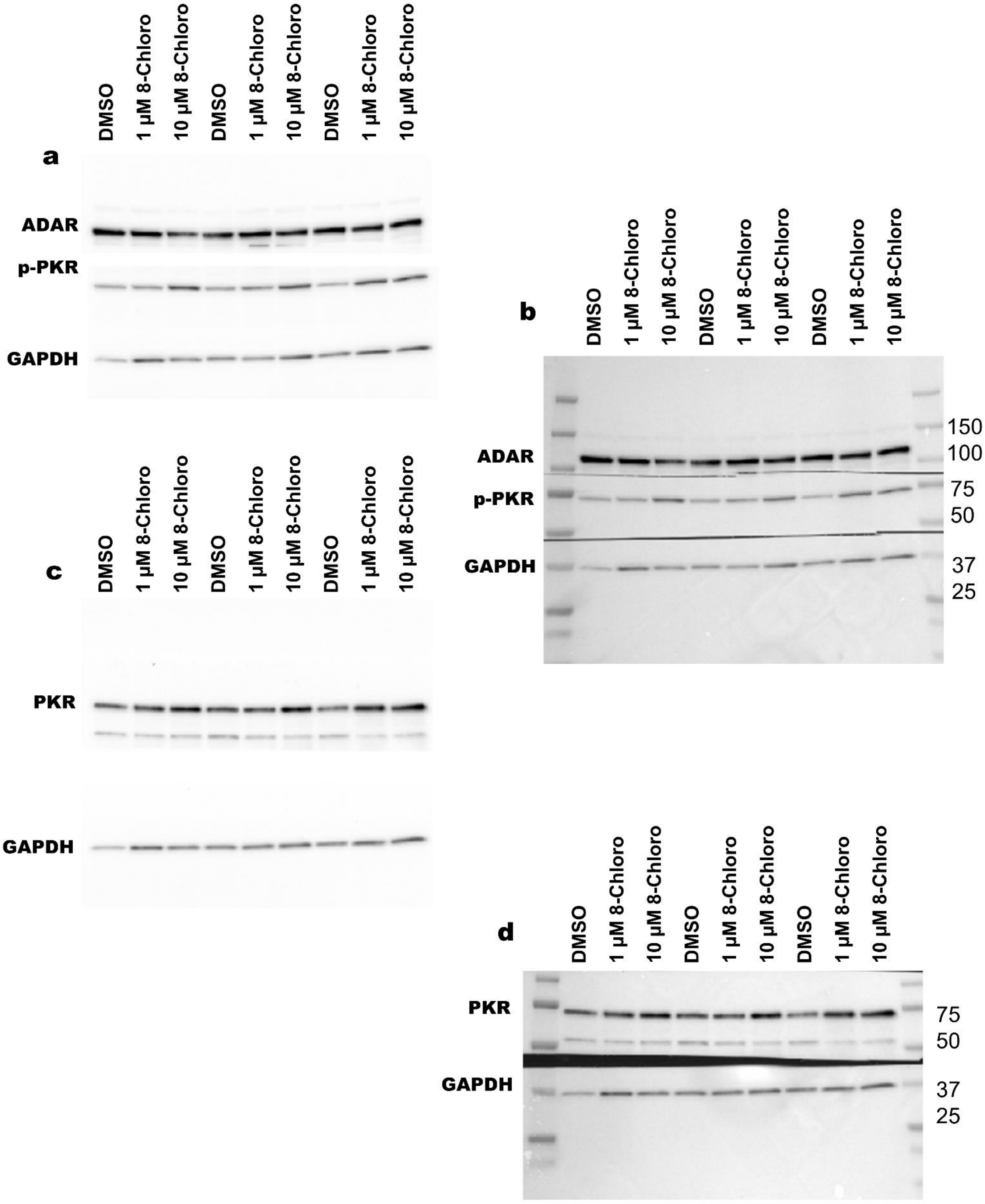


**Supplemental Figure 3:**

Uncropped immunoblots for HCC1806 treatment with 8-chloroadenosine (8-chloro) associated with Figure 3b, 3e and 3f. Panels **a** and **b** are the uncropped chemiluminescence images, panels **b** and **d** are the chemiluminescence images merged with colorimetric images to show the molecular weight marker.

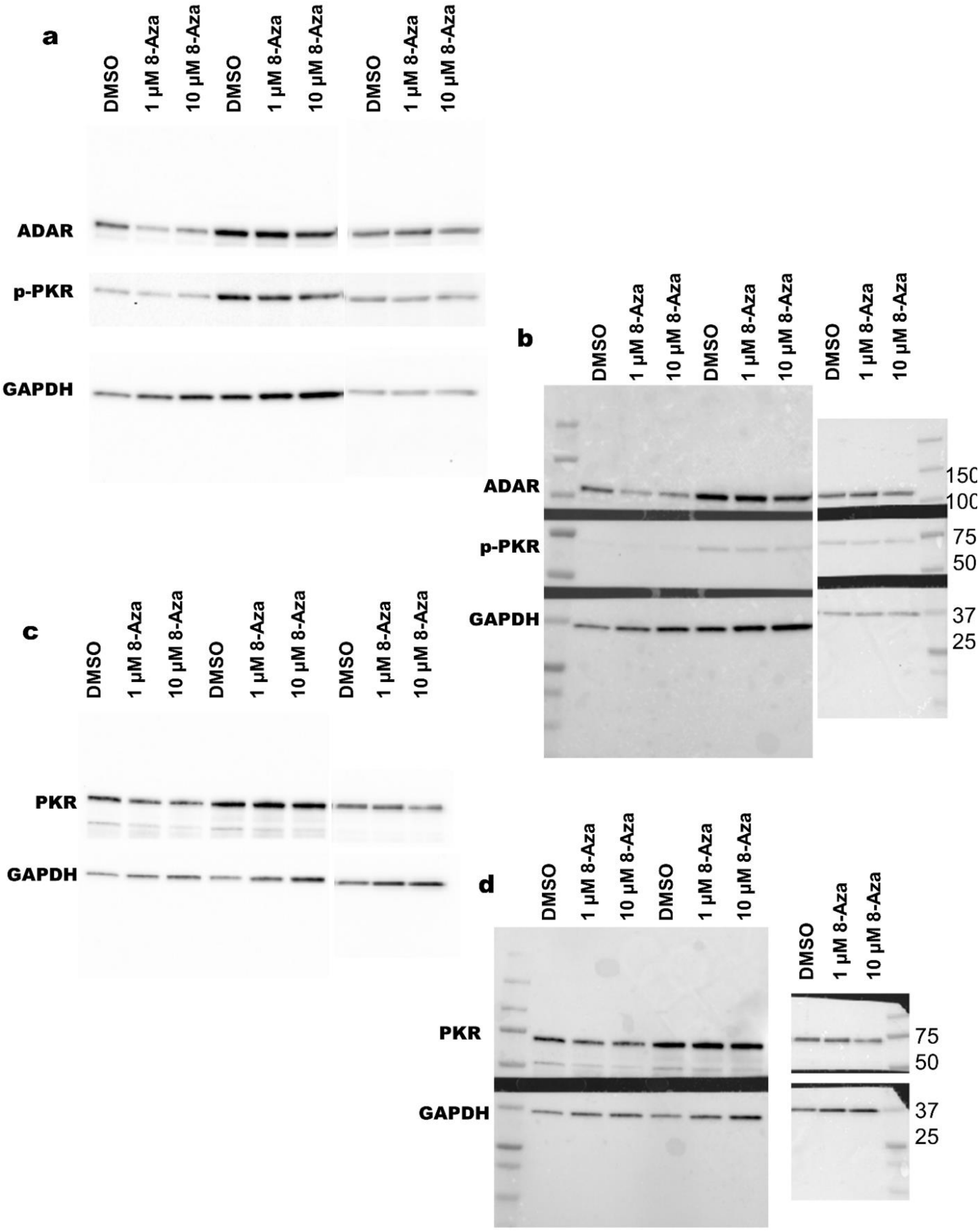


**Supplemental Figure 4:**

Uncropped immunoblots for MDA-MB-468 treatment with 8-azaadenosine (8-Aza) associated with Figure 3g-3i. Panels **a** and **b** are the uncropped chemiluminescence images, panels **b** and **d** are the chemiluminescence images merged with colorimetric images to show the molecular weight marker.

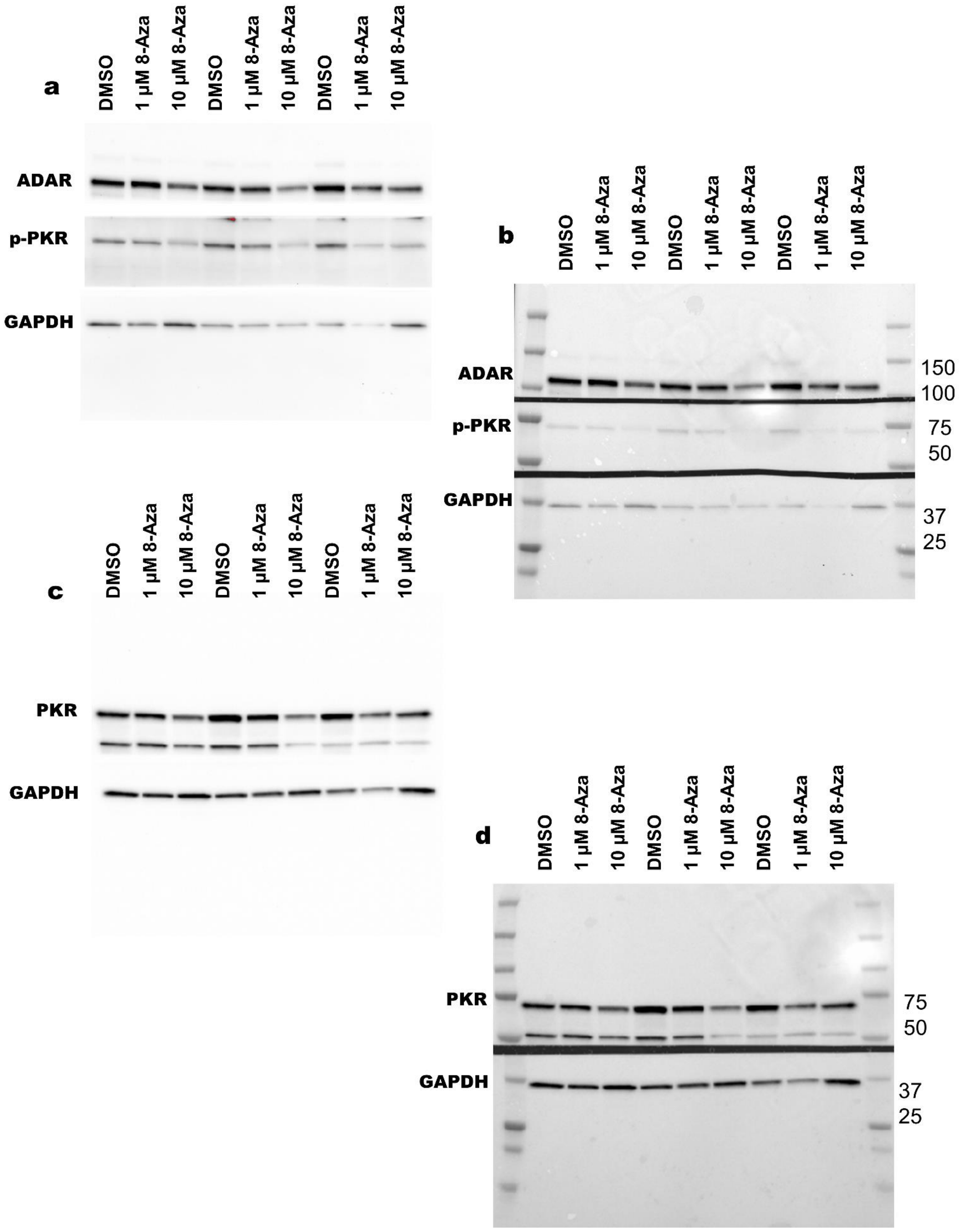


**Supplemental Figure 5:**

Uncropped immunoblots for HCC1806 treatment with 8-azaadenosine (8-Aza) associated with Figure 3g-3i. Panels **a** and **b** are the uncropped chemiluminescence images, panels **b** and **d** are the chemiluminescence images merged with colorimetric images to show the molecular weight marker.

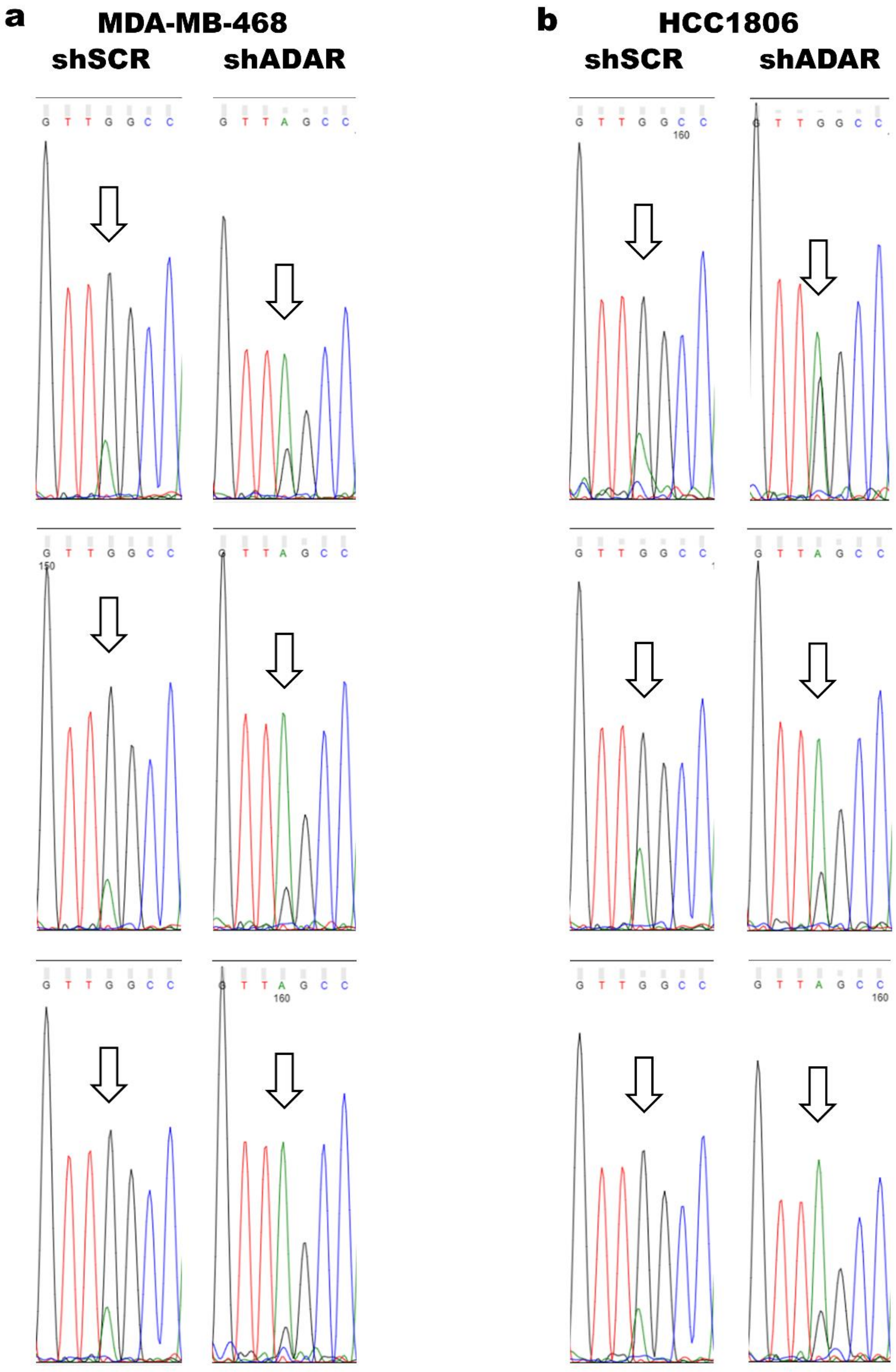


**Supplemental Figure 6:**

Chromatograms for all Sanger sequencing replicates associated with Figure 4a-b.

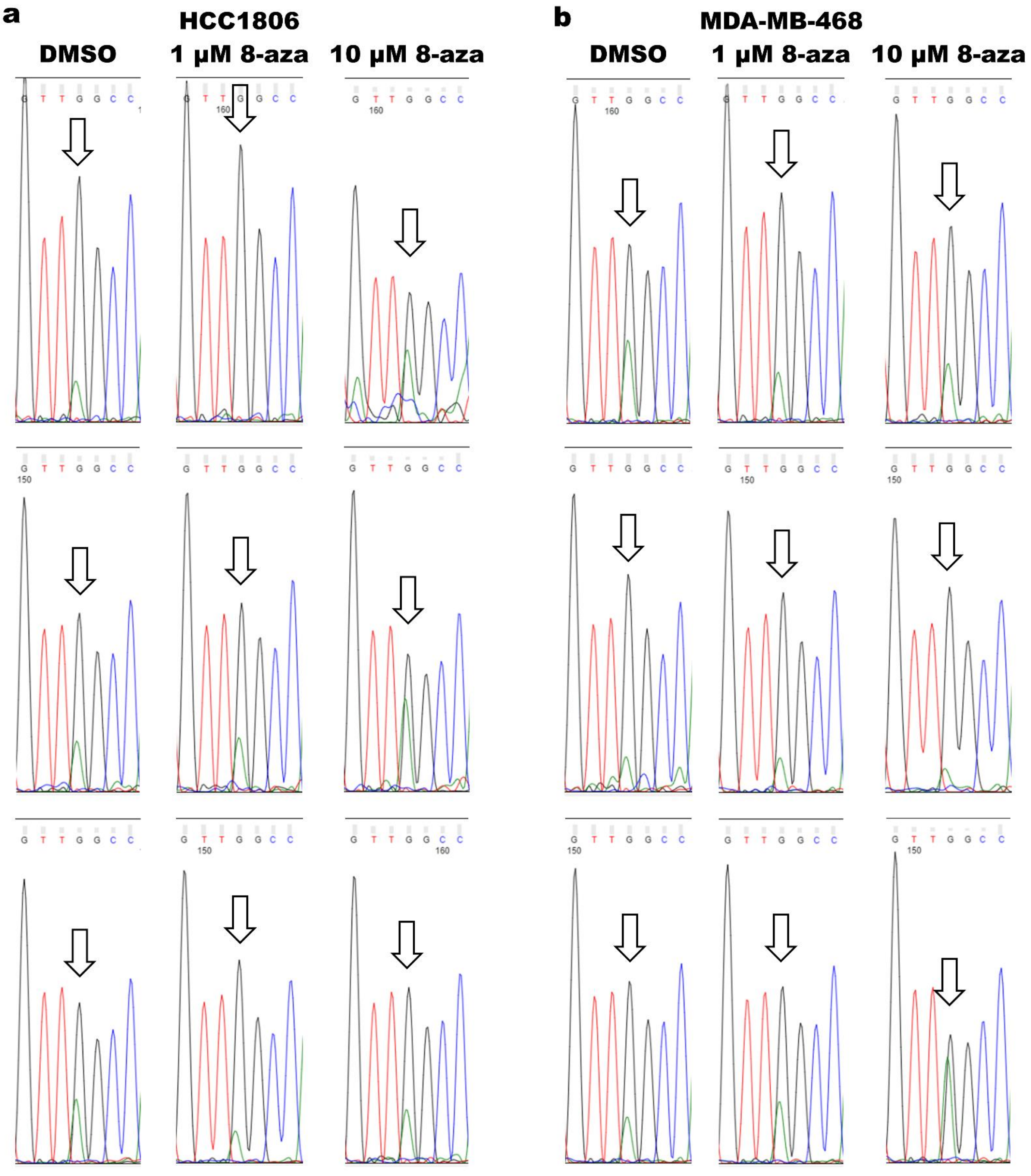


**Supplemental Figure 7:**

Chromatograms for all Sanger sequencing replicates associated with Figure 4c-d.

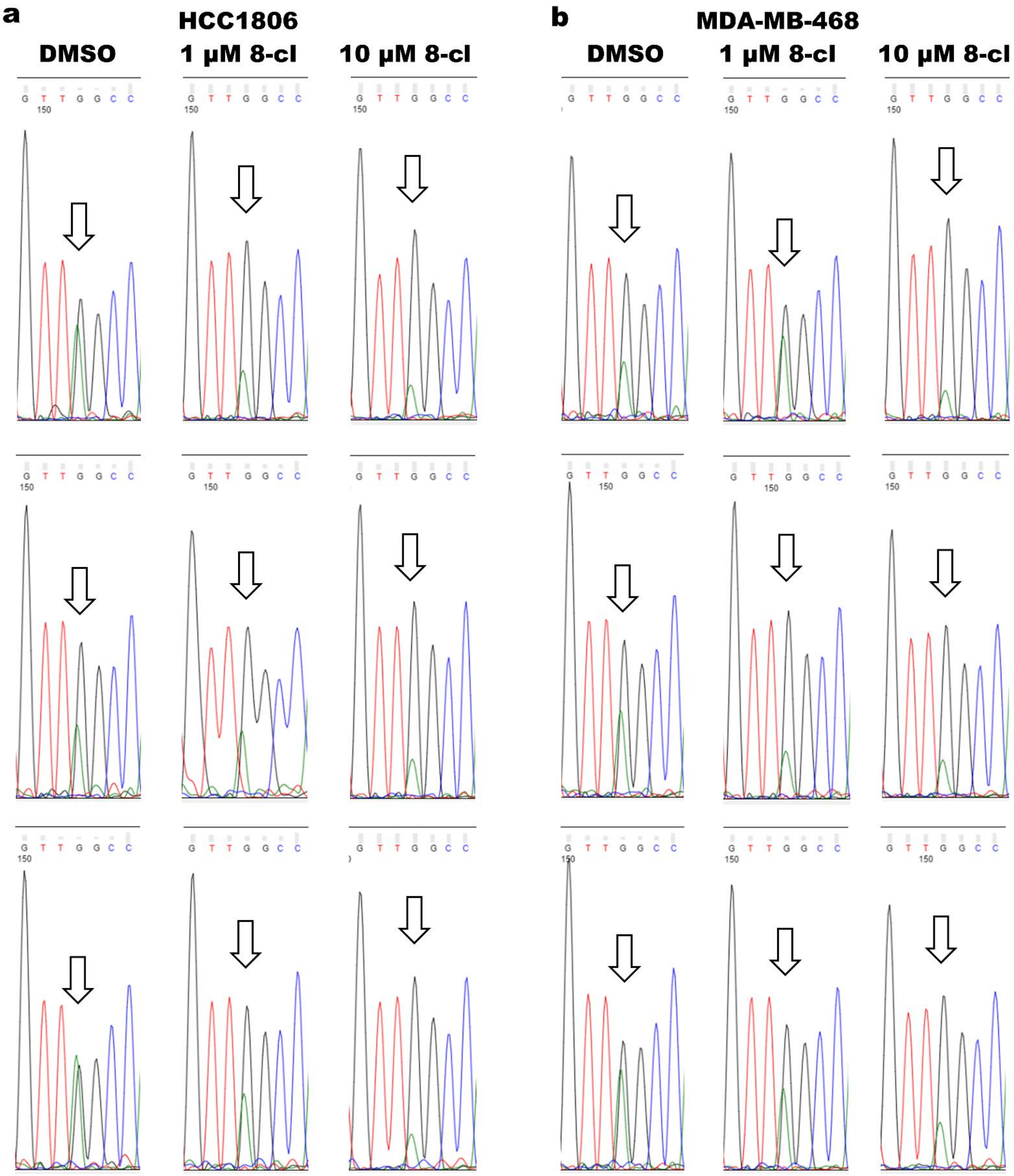


**Supplemental Figure 8:**

Chromatograms for all Sanger sequencing replicates associated with Figure 4e-f.
